# Non-uniform genetic effect sizes of variants associated with refractive error suggests gene-gene or gene-environment interactions are pervasive

**DOI:** 10.1101/483008

**Authors:** Alfred Pozarickij, Cathy Williams, Pirro Hysi, Jeremy A. Guggenheim, the UK Biobank Eye and Vision Consortium

## Abstract

Refractive error is a complex ocular trait controlled by genetic and environmental factors. Genome-wide association studies (GWAS) have identified approximately 150 genetic variants associated with refractive error. Among the known environmental factors, education, near-work and time spent outdoors have been demonstrated to have the strongest associations. Currently, the extent of gene-environment or gene-gene interactions in myopia is unknown. Here we show that the majority of genetic variants associated with refractive error show evidence of effect size heterogeneity, which is a hallmark feature of genetic interactions. Using conditional quantile regression, we observed that 88% of genetic variants associated with refractive error have at least nominally-significant non-uniform, non-linear profiles across the refractive error distribution. SNP effects tend to be strongest at the phenotype extremes and have weaker effects in emmetropes. A parsimonious explanation for these findings is that gene-environment or gene-gene interactions in refractive error are pervasive.

**Author summary:** The prevalence of myopia (nearsightedness) in the United States and East Asia has almost doubled in the past 30 years. Such a rapid rise in prevalence cannot be explained by genetics, which implies that environmental (lifestyle) risk factors play a major role. Nevertheless, diverse approaches have suggested that genetics is also important, and indeed approximately 150 distinct genetic risk loci for myopia have been discovered to date. One attractive explanation for the evidence implicating both genes and environment in myopia is gene-environment (GxE) interaction (a difference in genetic effect in individuals exposed to a high vs. low level of an environmental risk factor). Past studies aiming to discover GxE interactions in myopia have met with limited success, perhaps because information on lifestyle exposures during childhood has rarely been available. Here we used an agnostic approach that does not require information about specific lifestyle exposures in order to detect ‘signatures’ of GxE interaction. We found compelling evidence for widespread genetic interactions in myopia, with 88% of 150 known myopia genetic susceptibility loci showing an interaction signature. These findings suggest that GxE interactions in myopia are pervasive.

## Introduction

The prevalence of refractive error has risen steadily in the United States and other parts of the world in the past few decades [1, 2]. Nationally representative estimates for citizens aged 12-54 years derived from National Health and Nutrition Examination Surveys reported the prevalence of myopia as 41.6% in 1999-2004, compared to 25.0% in 1971-1972 [3]. By 2050 it is predicted that 49.8% of the world population will be myopic (4.8 billion individuals affected) [4]. Myopia is associated with axial elongation of the eye, which increases the risk of retinal detachment, myopic maculopathy, glaucoma, and other pathological complications, making it an increasingly common cause of visual impairment and blindness [5-7].

Susceptibility to myopia is determined both by genetic and environmental factors. Pedigrees in which high myopia segregates as a monogenic trait have pinpointed a number of genes whose normal function is required during refractive development [8-14]. The causative mutations responsible for these monogenic forms of high myopia are extremely rare in the population, and are characterized by profound, adverse effects on gene function. Genome-wide association studies (GWAS) have identified approximately 150 genetic variants associated with refractive error [15-18], including some overlap with monogenic disease gene loci [19]. These susceptibility variants are typically common in the population, yet individually have only small effects. Typically, each risk allele confers a shift towards a more negative (myopic) refractive error by a fraction of a diopter (D). In combination, however, the thousands of genetic risk variants thought to exist are estimated to explain at least 30% of the variation of refractive error [20-24]. The time children spend outdoors, time performing near-viewing tasks, and the number of years in education are also strongly associated with myopia development [25-32]. In randomized controlled trials, the incidence of myopia was significantly lower in children assigned to receive extra time outdoors during the school day [27-29]. Although associations between time spent performing near-viewing tasks and myopia have been reported less consistently than those for time outdoors, in the ALSPAC longitudinal birth cohort study, the amount of time children spent reading at the baseline age of 7 years-old explained approximately as much of the variation in refractive error over the next 8 years as the time the children spent outdoors at baseline [23]. In Mendelian randomization studies, each year in education has been estimated to cause a −0.25 to −0.40 D shift in refractive error [31, 32].

Conventionally, it is assumed that the ‘effect size’, i.e. the shift towards a more negative refractive error, of each myopia-predisposing genetic variant is consistent across the population. However, recent studies using the conditional quantile regression (CQR) technique to assess the consistency of effect sizes in phenotypes as diverse as BMI [33-35], height [33, 35], growth-rate of pigs [36], plant flowering time [37] and gene expression [38] have demonstrated that certain genetic variants exert differing effects depending on where an individual lies in the phenotypic distribution. For instance, using CQR, the BMI-susceptibility variant rs6235 in the *PCSK1* gene was recently shown to confer a shift towards a *lower* BMI in slim individuals and a *higher* BMI in obese individuals [35]. Such a non-constant effect size across quantiles of the phenotype distribution is a ‘signature’ of a variant involved in either a gene-gene (GxG) or gene-environment (GxE) interaction, or both. Crucially, unlike most other methods for detecting GxG and GxE interactions, the CQR approach does not require the identity of the environmental risk factor underlying a GxE effect to be pre-specified and measured in the study sample, nor the identity of the second genetic variant to be known when detecting GxG interactions. Instead, the presence of a likely GxG or GxE interaction can be evaluated using only genotype information for a genetic marker and phenotype information for the trait of interest (see Figure 1). Since GxE effects are implicated in susceptibility to myopia [39-42], and yet currently very few such interacting variants have been discovered, we aimed to comprehensively assess the known genetic variants associated with refractive error for involvement in interactions, using a CQR framework.

**Figure 1.**
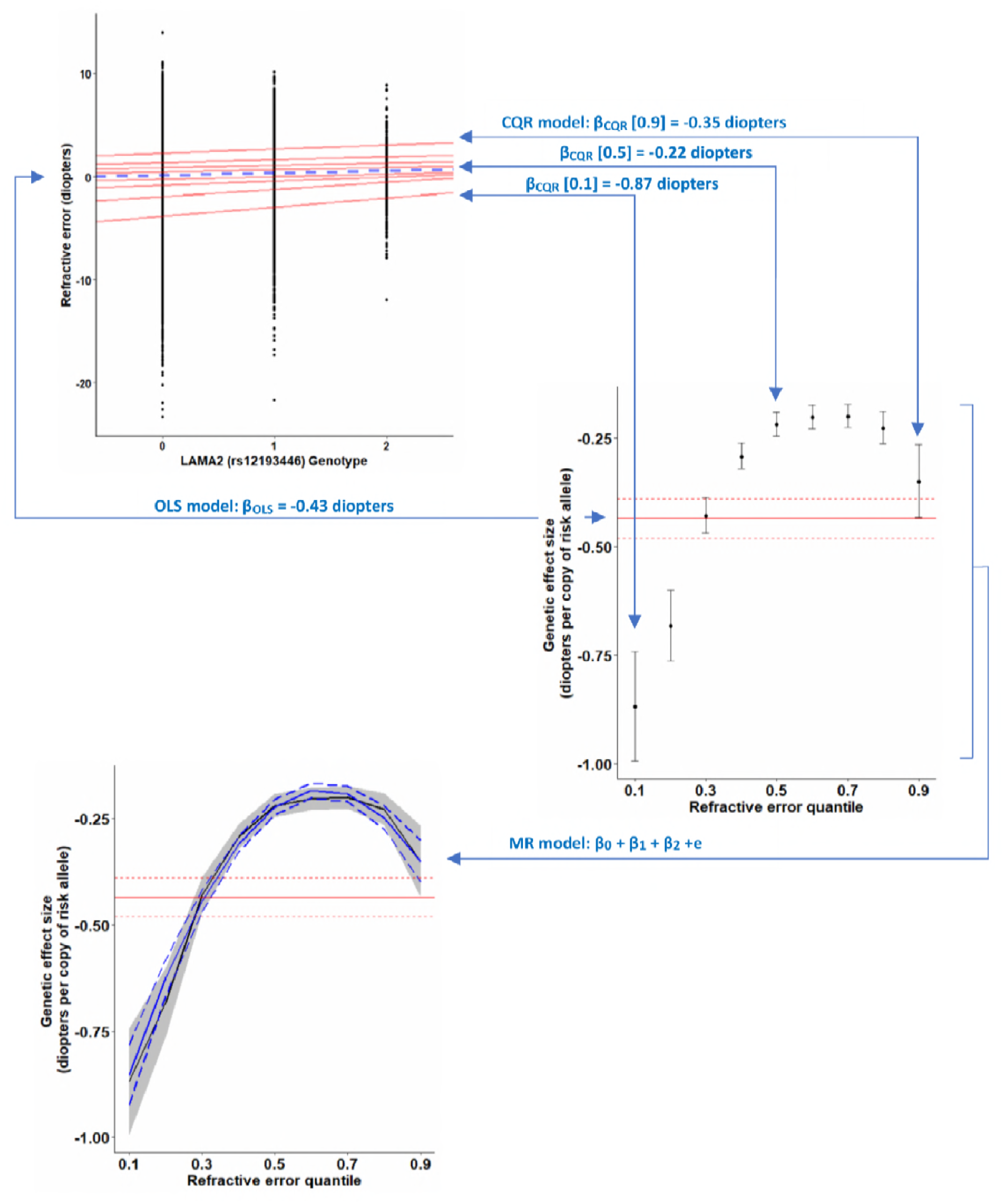
Conditional quantile regression (CQR) and meta-regression (MR) can identify if genetic effect size varies in individuals depending on their position in the trait distribution. In OLS, a SNP effect size is estimated under the assumption that it is the same for every person in the sample. In CQR, the SNP effect size is estimated at a specific quantile of the outcome distribution. The effect size may differ at different quantiles. After using CQR to estimate the SNP effect size at a range of quantiles, the uniformity of the SNP effect sizes is quantitatively assessed using MR.

## Results

### Type I error and power of CQR-MR

CQR is known to have a well-controlled type I error rate [43]. However, the *non-linear* meta-regression (MR) approach we used to combine results from across different quantiles has not been studied previously in the context of CQR to our knowledge. Therefore, we used the gold-standard method of permutation to examine how the type I error rate and power of the 3 terms of our CQR-MR model (β_0_, β_1_ and β_2_; i.e. the intercept, linear and quadratic terms) varied depending on the number of quantiles included in the MR model. As illustrated in Figure 2A & B, we evaluated MR models that included 19, 10, 9 or 5 quantiles from across the trait distribution. The main findings were: [1] there was a systematic inflation of the type I error rate for all 3 terms in the CQR-MR model (Figure 2C), [2] the type I error rate of CQR-MR was independent of MAF (Figure 2D), and [3] the model including 5 quantiles was overly conservative (Figure 2E). Fortunately, the systematic nature of this source of bias meant that it was straightforward to correct for (see *Methods* section).

**Figure 2.**
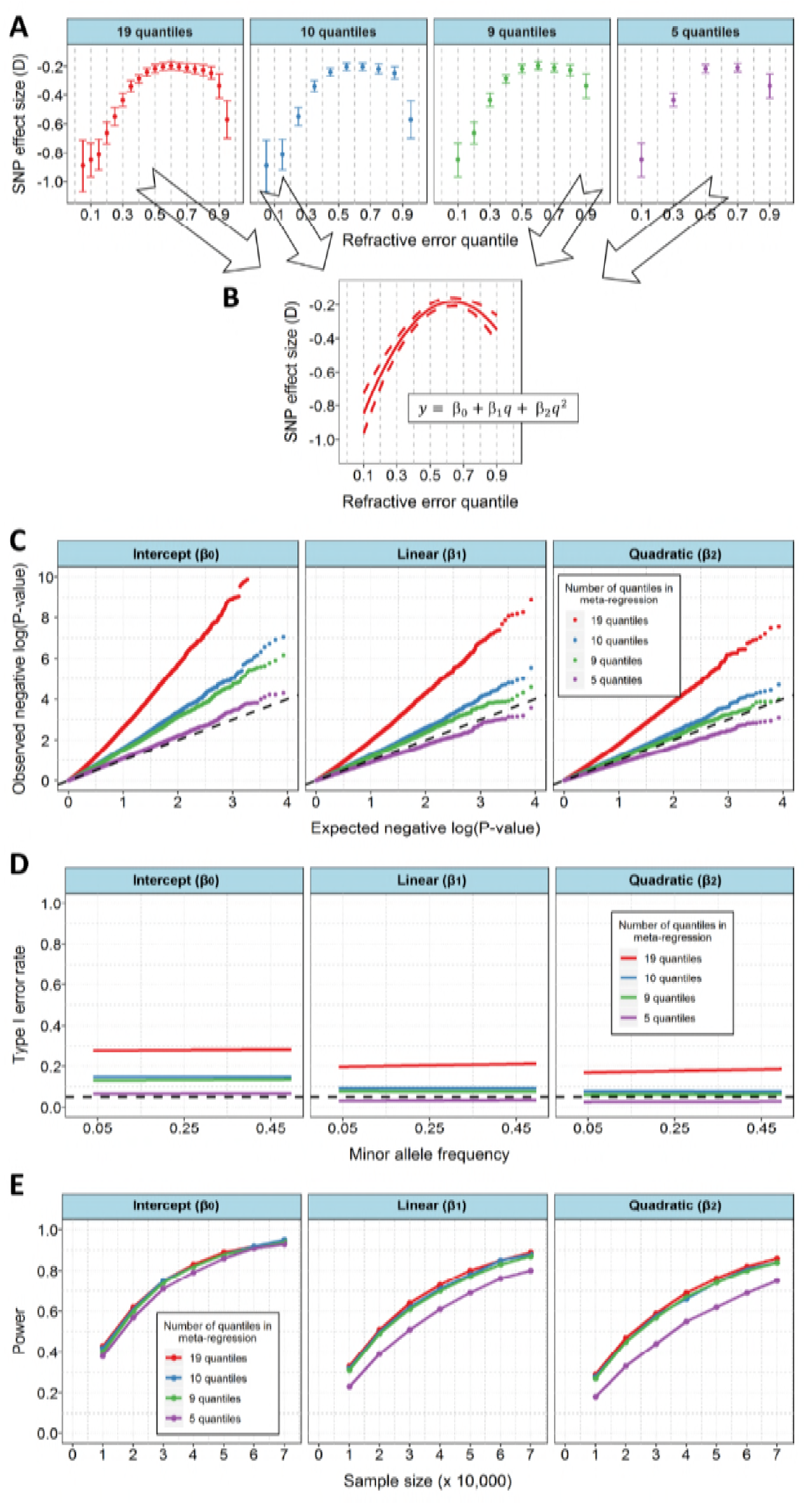
Type I error rate and power for MR models with different number of quantiles. **(A)** Illustration of genetic effect size estimates for CREAM variant rs12193446 from CQR carried out at 19, 10, 9 or 5 quantiles across the trait distribution. **(B)** MR was used to fit a quadratic function to the CQR results, in order to quantify the degree of non-uniformity and non-linearity of the CQR genetic effect size estimates. This yielded three coefficients describing the fit: intercept term β_0_, linear term β_1_, and quadratic term β_2_. **(C)** CQR-MR models were fit for 14,900 ‘null phenotype’ permutations. QQ-plots for observed versus expected p-values demonstrate systematic inflated test statistics (an excessive of low p-values) for MR models that included 19, 10 or 9 quantiles. The dashed black line is the line of unity. **(D)** The type I error rate for the models fit in (C). The dashed black line shows the correct type I error rate. Note the observed type I error rate is inflated for MR models including 19, 10 or 9 quantiles, and conservative for the 5 quantile MR model. **(E)** Relative statistical power of MR models including 19, 10, 9 or 5 quantiles, after adjusting for the inflated type I error rate. Note that power is lower for the 5 quantile MR model.

For the CQR-MR intercept term (β_0_), inflation of the type I error rate was apparent for all models, becoming progressively worse when greater numbers of quantiles were included in the MR. For example, the model with 19 quantiles had a type I error rate of approximately 0.30, while the model with 10 quantiles had a type I error rate of 0.16. The type I error rate for the model with 5 quantiles was 0.06, which was close to – but still above – the correct type I error rate (α= 0.05). The type I error rate for the CQR-MR linear and quadratic terms (β_1_ and β_2_ coefficients respectively) was also inflated for most of the CQR-MR models tested, with the degree of inflation again worsening when greater numbers of quantiles were included in the MR model (Figure 2C & D). However, for CQR-MR models that included only 5 quantiles, the type I error rate for the linear and quadratic terms was slightly conservative (approximately 0.04).

After correcting for the inflated type I error rate (see *Methods*), the statistical power was similar when either 19, 10 or 9 quantiles were included in the CQR-MR model, however power was reduced when only 5 quantiles were included in the MR model (Figure 2E).

In summary, CQR-MR models that included 19, 10 or 9 quantiles had inflated type I error rates for all 3 terms in the model, requiring the use of correction factors to account for this bias. An MR model that included only 5 quantiles had a conservative type I error rate, and was less powerful than MR models that included 9, 10 or 19 quantiles. Hence, for the analysis of CREAM variants, we used a CQR-MR model that included 9 quantiles. To correct the type I error rate, the intercept, linear and quadratic components of the model were adjusted using λ coefficients of 1.66, 1.23 and 1.10, respectively (i.e. observed Chi-squared statistics were divided by the relevant λ coefficient when calculating confidence intervals and p-values).

### Ordinary least squares (OLS) analysis of CREAM variants associated with refractive error

In the sample of 72,985 white British unrelated UK Biobank participants whose genotype data passed quality control and had phenotype information available, the average refractive error (avMSE) was −0.25 ± 2.67 D (mean ± SD) and the average age was 57.8 ±7.8 years. Of the 149 genetic variants associated with refractive error that were tested, reliable results could be obtained for 146 (for rs74764079, rs73730144 and rs17837871, with MAFs of 3%, 1% and 1%, respectively, there were fewer than 50 participants homozygous for the minor allele; hence these variants were excluded). After performing ordinary least squares (OLS) linear regression analysis to obtain SNP effects under the assumption of constant effect size across quantiles, the strongest effect was for the A allele of rs12193446 near the *LAMA2* gene, which was associated with a −0.43 D more negative refractive error (95% CI −0.48 to −0.39, p = 1.1 × 10^−77^), while the strongest association was observed for rs524952 (effect size = −0.26 D; p = 6.18 ×10^−79^; Table 1). The weakest effect was −0.03 D (95% CI −0.06 to +0.00, p = 0.05) for the T allele of *ZNF366* variant rs11952819. Full results are shown in Table S1. These effect sizes were extremely similar to those reported by the CREAM Consortium and 23andMe in a larger, but fully overlapping subset of UK Biobank participants that included related and unrelated individuals [18].

**Table 1.**
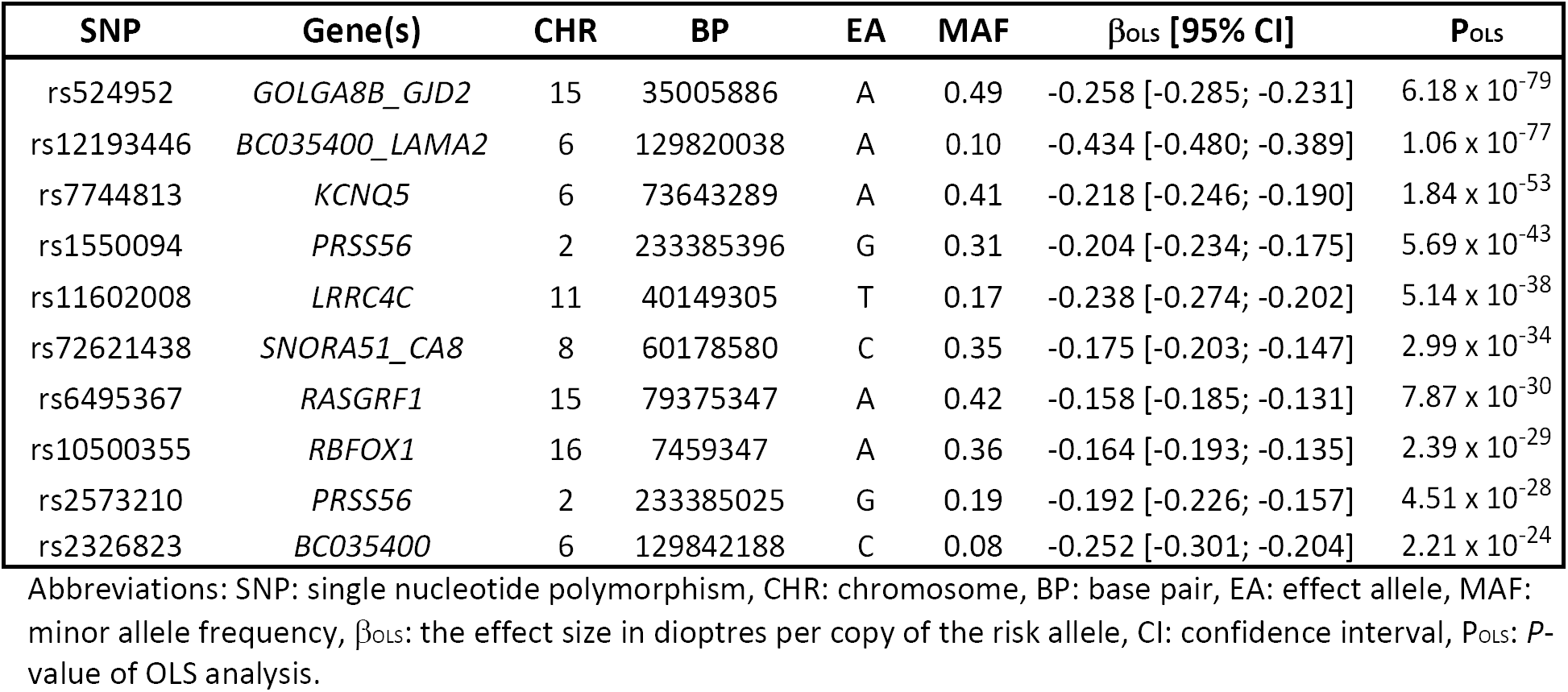
Summary statistics for 10 strongest associations with refractive error (avMSE) based on ‘conventional’ Ordinary Least Squares (OLS) regression.

### CQR-MR analysis of CREAM variants associated with refractive error

Conditional quantile regression was used to determine if effect sizes for the CREAM variants differed across quantiles of the refractive error distribution. A total of 146 variants were examined (the same set of variants examined above by OLS regression). As shown for a set of representative variants in Figure 3, we observed that most variants exhibited an inverse-U shaped effect size profile, with the strongest effect size at the extremes of the refractive error distribution and a minimum effect size near the 0.5 quantile (corresponding to emmetropic participants). Results for all variants are presented in Figure S1. A few variants, such as rs1649068 (*BICC1*) and rs9388766 (*L3MBTL3*) showed non-constant, yet more linear changes in effect size across quantiles of the refractive error distribution, thus yielding non-zero effects only for the extremes of the refractive error distribution. Finally, a small number of SNPs, such as rs9680365 (*GRIK1*) and rs7449443 (*FLJ16171-DRD1*), showed essentially flat effect size profiles that were similar to those obtained under the OLS assumption of a constant effect size across quantiles. Including principal components in the models did not change parameter estimates substantially, such that the same non-constant SNP effect distribution across the quantiles was observed. Variant rs12193446 near the *LAMA2* gene had the strongest effect across all the quantiles (Figure 2A & 3), reaching a maximum for moderate-to-highly myopic individuals at the 0.05 quantile: effect size = −0.9 D (95% CI −1.1 to −0.7).

**Figure 3.**
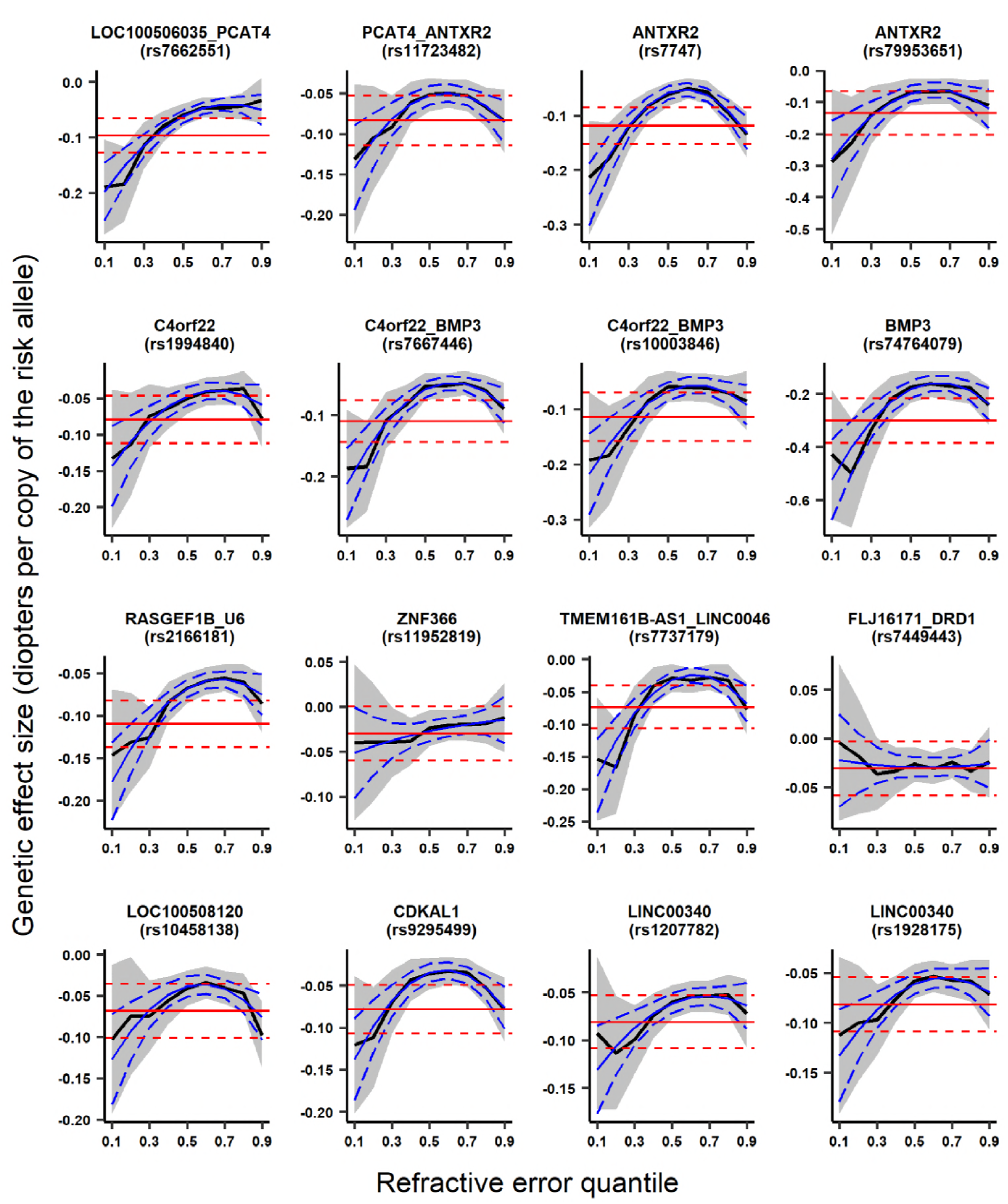
Changes in genetic effect size across the refractive error distribution for a representative subset of genetic variants associated with refractive error. Genetic effect size estimates from conditional quantile regression (CQR) are represented by the solid black line and their 95% confidence intervals are shown by the shaded grey region. Confidence intervals were calculated by using 10,000 Markov-chain-marginal-bootstrap replicates. The solid red line is the effect size estimate from conventional linear regression analysis with its 95% confidence intervals shown by the red dashed lines. Effect size estimates from meta-regression are shown with the solid blue line with corresponding 95% confidence intervals given by the dashed blue lines.

MR was used to quantitatively model the genetic effect size profile generated by quantile regression analysis. After correcting for the inflated type I error rate and accounting for multiple-testing by applying a Bonferroni adjusted p-value threshold of 0.05/(3 × 146) = 1.1 × 10^−4^, a total of 66 of the 146 (45%) CREAM variants had significant β_1_ (linear) or β_2_ (quadratic) meta-regression coefficients (Table 2 and Table S3). Thus, 45% of the genetic variants had differing effect sizes depending where in the refractive error distribution an individual lay, suggestive of the variant’s involvement in either a gene-gene or gene-environment interaction. The most extreme example was rs12193446 (*LAMA2*; Figure 2B & 3), which exhibited very strong evidence for a non-uniform effect size across quantiles (β_1_ component, p_adj_ = 2.12 ×10^−36^; β_2_ component, p_adj_ = 1.19 ×10^−30^). Of these 66 variants, 53 had significant β_2_ coefficients, indicating a non-linear effect size profile. Notably, only 18 of the 146 CREAM variants (12%) failed to show at least nominal evidence of an interaction effect (i.e. β_1_ component and β_2_ component, p < 0.05). A summary comparing these results for refractive error to those obtained by using height as a phenotype (and SNPs associated with height) is described in Supplementary Text S1, Tables S3 & S4, and Figure S2.

**Table 2.** Summary statistics for the 10 strongest associations with refractive error (avMSE) based on conditional quantile regression – meta-regression (CQR-MR). The intercept component (β_0_) CI and p-values have been adjusted using the ‘lambda GC approach’ by a factor of λ=1.66. The linear component (β_1_) CI and p-values have been adjusted by λ=1.22. The quadratic component (*β*_2_) CI and p-values have been adjusted by λ=1.10 to account for inflation of false positives in meta-regression.

| SNP | Gene(s) | $\beta_0$ [95% CI] | $P_{MR} - \beta_0$ | $\beta_1$ [95% CI] | $P_{MR} - \beta_1$ | $\beta_2$ [95% CI] | $P_{MR} - \beta_2$ |
| --- | --- | --- | --- | --- | --- | --- | --- |
| rs12193446 | BC035400_LAMA2 | -1.130 [-1.272; -0.988] | $8.07 \times 10^{-55}$ | 2.995 [2.529; 3.461] | $2.12 \times 10^{-36}$ | -2.363 [-2.765; -1.961] | $1.19 \times 10^{-30}$ |
| rs524952 | GOLGA8B_GJD2 | -0.673 [-0.758; -0.588] | $4.83 \times 10^{-54}$ | 1.797 [1.534; 2.06] | $7.47 \times 10^{-41}$ | -1.417 [-1.634; -1.200] | $1.68 \times 10^{-37}$ |
| rs7744813 | KCNQ5 | -0.543 [-0.631; -0.455] | $7.24 \times 10^{-34}$ | 1.402 [1.132; 1.672] | $2.15 \times 10^{-24}$ | -1.092 [-1.314; -0.870] | $5.75 \times 10^{-22}$ |
| rs11602008 | LRRC4C | -0.669 [-0.79; -0.548] | $2.60 \times 10^{-27}$ | 1.612 [1.250; 1.974] | $2.71 \times 10^{-18}$ | -1.131 [-1.421; -0.841] | $2.25 \times 10^{-14}$ |
| rs1550094 | PRSS56 | -0.521 [-0.624; -0.418] | $4.77 \times 10^{-23}$ | 1.441 [1.118; 1.764] | $2.08 \times 10^{-18}$ | -1.142 [-1.409; -0.875] | $4.90 \times 10^{-17}$ |
| rs72621438 | SNORA51_CA8 | -0.441 [-0.530; -0.352] | $2.06 \times 10^{-22}$ | 1.089 [0.817; 1.361] | $4.46 \times 10^{-15}$ | -0.821 [-1.044; -0.598] | $5.85 \times 10^{-13}$ |
| rs2326823 | BC035400 | -0.680 [-0.830; -0.530] | $6.17 \times 10^{-19}$ | 1.815 [1.341; 2.289] | $6.45 \times 10^{-14}$ | -1.429 [-1.831; -1.027] | $3.09 \times 10^{-12}$ |
| rs10500355 | RBFOX1 | -0.400 [-0.490; -0.310] | $3.63 \times 10^{-18}$ | 1.011 [0.734; 1.288] | $8.39 \times 10^{-13}$ | -0.775 [-1.003; -0.547] | $2.76 \times 10^{-11}$ |
| rs6495367 | RASGRF1 | -0.374 [-0.459; -0.289] | $7.17 \times 10^{-18}$ | 1.009 [0.747; 1.271] | $4.38 \times 10^{-14}$ | -0.833 [-1.049; -0.617] | $3.89 \times 10^{-14}$ |
| rs2573210 | PRSS56 | -0.501 [-0.621; -0.381] | $2.91 \times 10^{-16}$ | 1.414 [1.037; 1.791] | $1.94 \times 10^{-13}$ | -1.121 [-1.434; -0.808] | $2.26 \times 10^{-12}$ |

Abbreviations: SNP: single nucleotide polymorphism, CHR: chromosome, BP: base pair, EA: effect allele, $\beta_0$ : meta-regression intercept effect size in dioptres per copy of the risk allele, $\beta_1$ and $\beta_2$ : meta-regression coefficients for the linear and quadratic terms, respectively, CI: confidence interval.

### OLS and CQR-MR analysis of refractive error stratified by educational attainment

We derived a polygenic risk score (PRS) for refractive error derived from the 146 CREAM SNPs examined above. The effect size of the PRS was examined in participants categorized into 4 educational categories (the number of years spent in full-time education; *EduYears*). We estimated the effect of the PRS × *EduYears* interaction for *EduYears* groups 2, 3 and 4 with respect to the group with the least time spent in education (group 1; *EduYears* = 13-15 years). An OLS analysis yielded increasing evidence for a GxE interaction as *EduYears* increased: β_GxE_ = −0.05 D (95% CI −0.10 to 0.009; p = 0.1) for individuals in group 2 (*EduYears* = 16 years); β_GxE_ = −0.09 D (95% CI −0.15 to −0.03; p = 0.004) in group 3 (17-20 years); β_GxE_ = −0.17 D (95% CI −0.22 to −0.11; p = 7.53 ×10^−11^) in group 4 (21-26 years).

Quantile regression analysis revealed a highly non-uniform, non-linear relationship between the PRS effect size and refractive error quantile, which mirrored the inverted-U pattern observed for the majority of individual SNPs (Figure 4). Notably, the PRS effect size differed across educational attainment strata. For participants from the myopic tail of the refractive error distribution (i.e. quantiles <0.2), more time spent in education was associated with a larger PRS effect size (Figure 4). For example for those in the 0.1st quantile, a 1 SD increase in PRS was associated with a −0.82 D (95% CI −0.90 to −0.73; p = 8.9 × 10-83) more negative refractive error in the lowest educational stratum, yet a −1.11 D (95% CI −1.18 to −1.02; p = 1.17 ×10^−155^) more negative refractive error for those in the highest education stratum. The largest change in PRS effect size due to such an interaction with education was 0.57 D at the 0.2^nd^ quantile. The PRS effect size difference associated with *EduYears* group was smallest in emmetropes: For instance, the PRS effect size was within a narrow range of −0.25 to −0.37 D for participants in the 0.6 quantile, irrespective of their level of education. For participants in the hyperopic tail of the refractive error distribution (quantiles >0.8), the PRS effect size was smaller in those with greater educational attainment, opposite to the relationship seen in the myopic tail. Thus, for example, for hyperopic participants in the 0.9 quantile, a 1 SD *reduction* in PRS was associated with a +0.62 D (95% CI +0.69 to +0.55; p = 6.55 × 10^−68^) effect on refractive error in those in the lowest education stratum (*EduYears* group 1), yet only a +0.41 D (95% CI +0.44 to +0.38; p = 9.53 ×10^−193^) effect in those from the highest education stratum (*EduYears* group 4).

**Figure 4.**
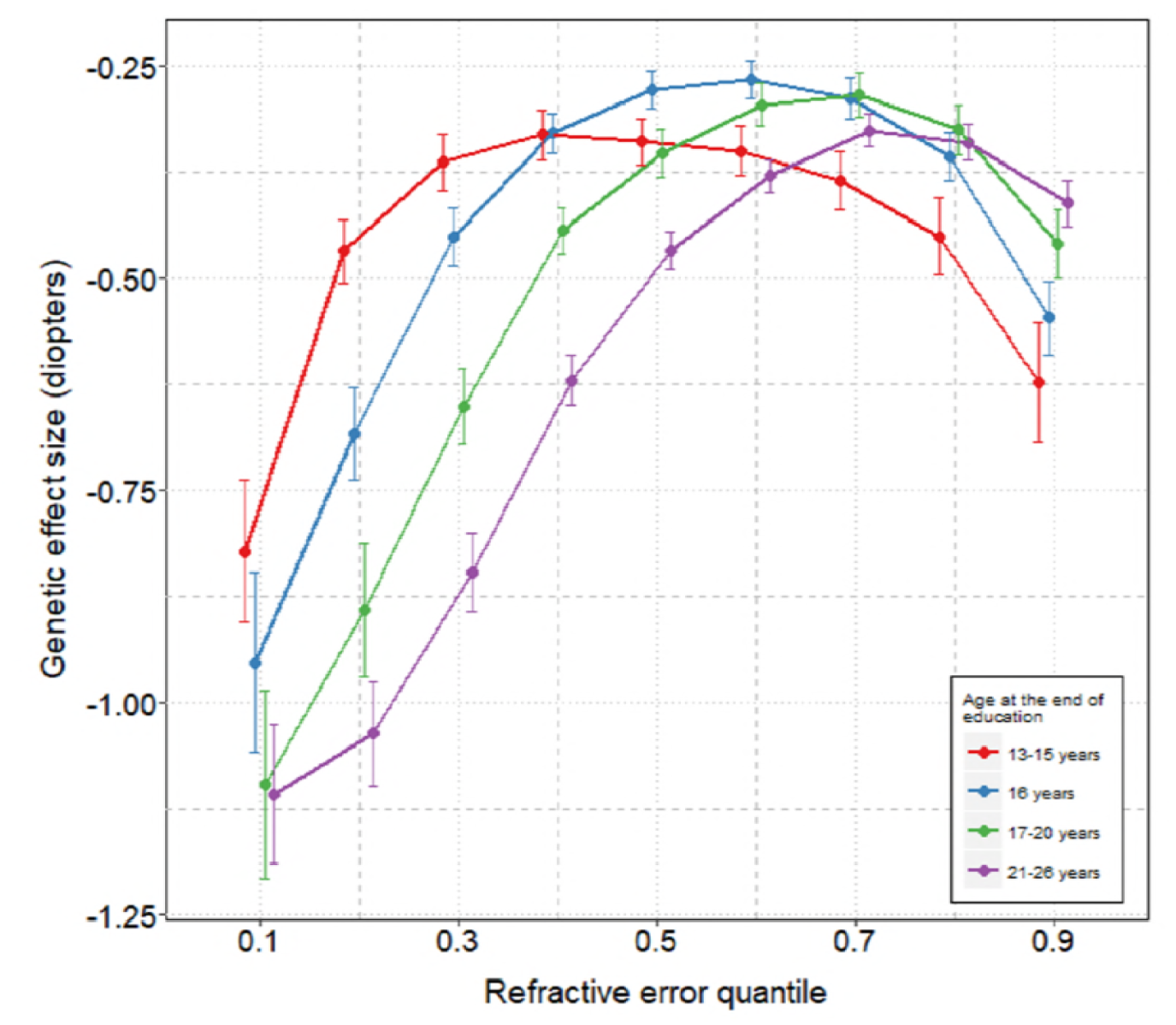
The effect of educational attainment (*EduYears*) on refractive error varies across quantiles of the refractive error distribution. Each line represents the PRS effect size across quantiles for individuals with different times spent in education. Error bars show 95% confidence intervals.

## Discussion

The detection of non-uniform genetic effect sizes across quantiles of the trait distribution by CQR-MR has previously been proposed as a method for identifying SNPs involved in GxE or GxG interactions, without any prior knowledge of the identity of the interacting variant or environmental exposure [35]. Here, we studied changes in genetic effect size across the refractive error trait distribution in SNPs previously shown to be associated with this trait, and found evidence for non-uniform effects in 88% of the 146 variants tested. This suggests that the majority of genetic variants associated with refractive error interact with environmental risk factors or other genetic variants to exert their effects. As an exemplar environmental exposure, years spent in education was shown to have a profound influence on the risk of myopia conferred by a PRS for refractive error. The interaction effect for education was greatest in individuals with moderate-to-high myopia, in whom the PRS was associated with a more than 0.50 D difference in refractive error between those in the lowest vs. highest education group.

Under the assumption that no gene-environment or gene-gene interaction modifies the effect of genetic variant, we would expect the distribution of SNP effects to be constant across all quantiles. Using quantile regression, we found that 45% of SNPs known to be associated with refractive error showed compelling evidence of a non-uniform effect size. Variants typically had inverse-U shaped effect size profiles, with the strongest effects observed at the phenotype extremes, and effects closer to zero in emmetropes. Very few SNPs had constant effects across all quantiles of the sample distribution that matched those assumed to occur when fitting conventional OLS models. An appealing explanation for these findings is the process of “emmetropization”, in which the rate of axial eye elongation during infancy is fine-tuned by a visual feedback loop in order to maintain a sharp retinal image [44]. Emmetropization may therefore act as a buffer against the myopia- or hyperopia-predisposing effects of risk variants.

Because refractive error is known to be influenced by multiple environmental factors [23, 23-32] and there is prior evidence of GxE interaction effects impacting refractive development – yet, to date, no evidence of GxG interaction – we suggest that the most parsimonious explanation for our quantile regression results is that GxE interactions are commonplace during myopia development. Prior to this work, only a handful of specific GxE interactions have been revealed for refractive error [39-42], predominantly in samples from south-east Asia in which the prevalence of myopia is extremely high. These included SNP × education interactions involving rs2969180 (*SHISA6-DNAH9*) (−0.28 D, p = 4.78 × 10^−4^), rs524952 (*GJD2*) (−0.23 D, p = 3.8 × 10^−3^), rs2137277 (*ZMAT4-SFRP*) (−0.42 D, p = 2.1 × 10^−3^), rs12511037 (*AREG*) (−0.89 D, p = 6.87 × 10^−11^), rs13215566 (*GABRR1*) (−0.56 D, p = 8.48 × 10^−5^), and rs12206610 (*PDE10A*) (−0.72 D, p = 2.32 ×10^−8^); all with effect sizes estimated using OLS models [41, 42]. One gene-environment interaction involving rs188663068 (*APLP2*) and time spent reading in childhood (−0.6 D, p = 7.1 × 10^−4^) was identified in a British population and was also identified to be responsible for environmentally induced myopia in a transgenic mouse model [40]

Repeating our CQR-MR analyses for the variants most strongly associated with height identified by the GIANT consortium [45] revealed scant evidence for variants with non-uniform effect sizes across quantiles. This finding is consistent with previously-observed results for this trait [35, 46]. Since height is known to have a very high narrow-sense heritability [45], our findings point to a lack of GxE or GxG interactions for height SNPs that have strong marginal associations.

This study has several limitations. First, as with methods that test for a difference in variance between different genotype groups (“variance heterogeneity” analysis) [47-50], the CQR-MR framework aims to detect heteroscedasticity in SNP effects, i.e. when SNP effects are not constant across the phenotype distribution, this is an indication for potential involvement of interacting factors. However, mechanisms other than GxE or GxG interactions can also result in variance heterogeneity. These include, a) parent-of-origin effects [48], where genotypic variance in heterozygous individuals depends on which parent an allele was transmitted from and b) allelic heterogeneity [45, 51, 52], where multiple genotypes in linkage disequilibrium influence the same phenotype. Second, our CQR study of PRS-education interaction effects was restricted to testing a PRS rather than individual SNPs, because of the limited sample size. Therefore, our results were unable to highlight which specific SNPs interact with education. Third, our permutation analyses showed that CQR-MR models can have inflated type I error rates, and yield high numbers of false positives. This necessitated the use of a correction factor to adjust standard errors and p-values, making CQR-MR analyses more complex than standard OLS methods. The inflated type I error rate of CQR-MR is likely due to two reasons: the non-normal distribution of refractive error (we found the type I error rate was lower for CQR-MR analysis of height; Figure S3) and the fact that meta-regression assumes that SNP effects across quantiles are estimated independently, whereas they were in fact estimated for the same sample.

In summary, our study provides evidence that most genetic variants currently-known to be associated with susceptibility to refractive error have non-uniform effects across refractive error quantiles. The most parsimonious explanation is that these variants are subject to interaction effects, involving either additional gene variants or environmental risk factors. These effects typically have major impacts, such that effect sizes are often 2-to-3-fold greater for individuals in the phenotype extremes compared to those in the centre. This variation in effects remains hidden when conventional OLS regression methods are used to detect main and interaction effects. Thus, the complex interplay between genetic and environmental factors uncovered in this work explains some of the missing heritability for refractive error.

## Methods

### Study Participants and Quality Control

*UK Biobank*. The UK Biobank project is an ongoing cohort study of approximately 500,000 UK adults aged 40 to 70 years-old when recruited (2006-2010) [53]. Ethical approval for the study was obtained from the National Health Service National Research Ethics Service (Ref 11/NW/0382) and all participants provided written informed consent. Participants provided a blood sample, from which DNA was extracted and genotyped using either the UK BiLEVE Axiom array or the UK Biobank Axiom Array [54]. We analysed data from the July 2016 data release for genetic variants in 488,377 individuals imputed to the HRC [55] reference panel.

Participants self-reported whether they had a university or college degree. An ophthalmic assessment was introduced towards the latter stages of UK Biobank recruitment, hence only about 25% of participants were examined. Refractive error was measured using non-cycloplegic autorefraction (Tomey RC5000; Tomey GmbH Europe, Erlangen-Tennenlohe, Germany). The mean spherical equivalent (MSE) refractive error was calculated as the sphere power plus half the cylinder power, and averaged between the two eyes (avMSE). Individuals who self-reported any of the following eye disorders were excluded from the analyses: cataracts, “serious eye problems”, “eye trauma”, a history of cataract surgery, corneal graft surgery, laser eye surgery, or other eye surgery in the past 4 weeks. Individuals whose hospital records (ICD10 codes) indicated a history of the following were also excluded: cataract surgery, eye surgery, retinal surgery, or retinal detachment surgery. Of 488,377 individuals with genetic information, samples were excluded due to: Ocular history (n=48,145, see above), withdrawal of consent (n=8), self-report of non white-British ethnicity or genetic principal components indicative of non-European ancestry (n=69,938), outlying level of genetic heterozygosity (n=648), or refractive error not measured (n=283,352). The remaining 86,286 individuals were tested for relatedness using the --rel-cutoff command in PLINK v2 [56]. A genetic relationship matrix was created using a linkage disequilibrium (LD)-pruned set of well-imputed variants (with IMPUTE2 r^2^ > 0.9, minor allele frequency (MAF) > 0.005, missing rate ≤ 0.01, and ‘rs’ variant ID prefix). LD-pruning was accomplished by using the --indep-pairwise 50 5 0.1 command in PLINK v2 [56]. One member of each pair with genomic relatedness greater than 0.025 was excluded. This resulted in a final sample size of 72,985 unrelated individuals of European ancestry.

### Selection of Genetic Variants

#### Variants associated with refractive error

We assessed 149 genetic variants that showed genome-wide (marginal) association (p < 5 ×10^−8^) with refractive error in a recent meta-analysis carried out by the CREAM Consortium and 23andMe and that showed evidence of independent replication in the UK Biobank sample [18]. We coded the risk allele as the allele associated with a more negative refractive error.

#### Variants associated with height

For comparison, we also examined genetic variants associated with height. GWAS summary statistics were obtained from Wood et al. [45]. We restricted our analyses to the 149 genetic variants with the strongest association (i.e. those with the lowest p-values).

### Statistical Analysis

The ‘conventional’ effect size of each of the 149 refractive error genetic variants (i.e. under the assumption of a constant effect size across the full sample) was estimated using a linear regression model, with refractive error averaged between the 2 eyes (avMSE) as the dependent variable and genotype, age, age-squared, sex and a binary variable indicating genotyping array fitted as covariates. Conditional quantile regression [43] was carried out using the *quantreg* package in R, using the same set of covariates as above. We used 10,000 Markov-chain-marginal-bootstrap replicates to calculate standard errors. As a sensitivity analysis, we also tested linear regression and quantile regression models with the first 10 principal components included as covariates, to determine if confounding with residual ancestry influenced the results.

SNP effect estimates and their standard errors from quantile regression at 9 different quantiles (0.1, 0.2, 0.3, …, 0.9) were meta-regressed using a mixed-effects model (*metafor* package in R [57]) with the estimated SNP effect at each quantile modelled as the dependent variable and the quantile at which these estimates were obtained as the independent variable. A term for quantile-squared was also included in the meta-regression model to test for non-linear genetic effects across quantiles, resulting in the model: y = β_0_ + β_1_*q* + β_2_*q*^2^ + *e* (where, β_0_ is an intercept term, β_1_ and β_2_ are coefficients describing the linear and quadratic change in SNP effect across quantiles of the trait distribution, respectively, *q* are the quantiles, and *e* is the error term). Figure 1 illustrates the conditional quantile regression and meta-regression model fitting strategy.

### Permutation-based assessment of type I error rate and power

To assess the performance of our CQR-MR analysis model, its type I error and power were evaluated using the gold-standard method of permutation. The type 1 error rate was assessed in two ways. Firstly, we simulated genotypes for ‘null’ SNPs and tested for an association between the true phenotype and the null SNP genotype. Secondly, we permuted phenotype values amongst individuals in the sample, and tested for an association between the null phenotype and the observed (true) SNP genotypes.

#### Null phenotype

The avMSE phenotype of the 72,985 individuals in the analysis sample was permuted 100 times. For each permutation, the association between the null phenotype and the genotype of each of the 149 CREAM SNPs was assessed using CQR-MR. The type 1 error rate was calculated as the proportion of SNPs with P<0.05 for each of the three meta-regression coefficients (β_0_, β_1_ and β_2_) from the total of (100 × 149) 14,900 permutations. *Null SNPs*. The 72,985 individuals in our analysis sample were independently assigned genotypes for a biallelic SNP with MAF ranging from 0.05 to 0.45, simulated from a binomial distribution. Association between avMSE and the genotype of the null SNP was assessed using CQR-MR. The type 1 error rate was calculated as the proportion of SNPs with P<0.05 for each of the three meta-regression coefficients (β_0_, β_1_ and β_2_) after simulating 10,000 null SNPs.

To obtain a relative indication of statistical power, the 149 CREAM SNPs were tested for association with the observed avMSE refractive error phenotype in samples of varying size. Specifically, from the full sample of 72,985 individuals, we selected a random sample of 10,000 to 70,000 individuals, in steps of 10,000, and tested each of the 149 CREAM SNPs for association. This procedure was repeated 20 times. Power was computed as the proportion of replicates in which the null hypothesis was rejected at a nominal significance level of *α* = 0.05 (i.e. under the assumption that all 149 SNPs truly had non-linear effect sizes across quantiles). The total number of tests used for these power evaluations was 149 × 7 ×20 = 20,860. The same set of covariates as in original analysis was included in the CQR step when assessing power and type 1 error.

As described above, our ‘standard’ CQR-MR analysis model consisted of carrying out quantile regression at 9 different quantiles (*q* = 0.1 to 0.9 in steps of 0.1) followed by meta-regression of the resulting genetic effect size estimates. To explore the effect of selecting more or fewer than 9 quantiles, we also assessed the type 1 error rate and power when meta-regression was carried out with: a) 19 quantiles, *q* = 0.05 to 0.95 in steps of 0.05; b) 10 quantiles, *q* = 0.05 to 0.95 in steps of 0.1; c) 5 quantiles, *q* = 0.1 to 0.9 in steps of 0.2. For simplicity, we refer to these CQR-MR models by the number of quantiles included in the meta-regression, i.e. 19, 10, 9, or 5; with 9 quantiles being our ‘standard’ approach (Figure 2A & B).

### Adjustment for the inflated type 1 error rate

Meta-regression was found to produce a systematically inflated type 1 error rate (see *Results*). To adjust for this source of bias, we calculated ‘inflation factors’ (λ_β0_, λ _β1_ and λ _β2_) analogous to the use of λ_GC_ for genomic control [58], using the results from the ‘null phenotype’ permutation analyses described above. *P*-values and confidence intervals for each term (β_0_, (β_1_ and (β_2_) in the meta-regression were adjusted by their respective inflation factor with the equation: 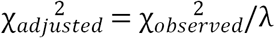, where λ was calculated as the adjusted observed observed median chi-squared statistic from the ‘null phenotype’ permutations divided by the expected median chi-squared statistic with 1 df. Noting that Z= (β/se and χ^2^ = Z^2^, meta-regression confidence intervals were calculated by adjusting standard errors: seadjusted 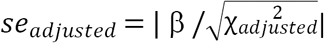

### Gene-Environment interaction with education

To test for the presence of gene-environment interaction, we constructed a polygenic risk score (PRS) by counting the number of risk alleles (0, 1 or 2) carried by each individual. We did not weight these by SNP effect sizes in order to avoid introducing bias by using weights obtained from, and applied in, the same sample (UK Biobank). ‘Age completed full-time education’ (*EduYears*) was selected as an exemplar environmental variable. UK Biobank participants with a university degree were not asked the age they completed full-time education, hence these individuals were assumed to have completed their education at the age of 21 years. Age completed education categories with low counts were merged with adjacent categories, resulting in 4 final *EduYears* categories: 13-15, 16, 17-20 and 21-26 years. Firstly, we fitted the following model using linear regression, avMSE = PRS + EduYears + PRS×EduYears + age + age^2 + sex + genotyping array. This was used to estimate the regression coefficients for the PRS×EduYears (β_GxE_) term under the assumption of constant effect size across quantiles. As an alternative, we carried out a CQR-MR analysis stratified by *EduYears* category. Table 3 provides a summary of refractive error distribution for each *EduYears* category, prior to merging groups with low numbers of participants.

**Table 3.** Distribution of refractive error per education attainment category.

| **Age at the end of education** | 13 | 14 | 15 | 16 | 17 | 18 | 19 | 20 | 21 | 22 | 23 | 24 | 25 | 26 |
| --- | --- | --- | --- | --- | --- | --- | --- | --- | --- | --- | --- | --- | --- | --- |
| **Refractive error in diopters mean (SD)** | 0.35<br>(2.71) | 0.71<br>(2.34) | 0.72<br>(2.28) | -0.004<br>(2.43) | -0.16<br>(2.53) | -0.53<br>(2.66) | -0.52<br>(2.62) | -0.59<br>(2.58) | -0.78<br>(2.85) | -0.51<br>(2.71) | -0.27<br>(2.57) | -0.21<br>(2.85) | -0.004<br>(2.04) | 0.08<br>(2.43) |
| **N** | 419 | 411 | 12,274 | 15,789 | 5623 | 6193 | 1453 | 897 | 27,897 | 680 | 317 | 138 | 152 | 293 |

## Description of Supplemental Data

Supplemental Data include an additional 3 figures, 4 tables, and 1 text file.

**Text S1**. CQR-MR analysis of GIANT consortium variants associated with height

**Table S1**. Summary statistics for ‘conventional’ Ordinary Least Squares (OLS) regression size estimates for association with refractive error (avMSE).

**Table S2**. Summary statistics for CQR-MR effect estimates for SNPs associated with refractive error (avMSE).

**Table S3**. Summary statistics for ‘conventional’ Ordinary Least Squares (OLS) regression size estimates for association with height.

**Table S4**. Summary statistics for CQR-MR effect estimates for SNPs associated with height.

**Figure S1**. Changes in genetic effect size across the refractive error distribution for genetic variants associated with refractive error.

**Figure S2**. Changes in genetic effect size across the height distribution for a selected subset of genetic variants associated with height.

**Figure S3**. Sample distributions for refractive error and height.

## Acknowledgments

This research has been conducted using the UK Biobank Resource (applications #17351). We thank Arkan Abadi and David Meyre (McMaster University) for advice on R codes. The work was funded by the National Eye Research Centre grant SAC015 (JAG, CW), and an NIHR Senior Research Fellowship award SRF-2015-08-005 (CW). UK Biobank was established by the Wellcome Trust; the UK Medical Research Council; the Department for Health (London, UK); Scottish Government (Edinburgh, UK); and the Northwest Regional Development Agency (Warrington, UK). It also received funding from the Welsh Assembly Government (Cardiff, UK); the British Heart Foundation; and Diabetes UK. Collection of eye and vision data was supported by The Department for Health through an award made by the NIHR to the Biomedical Research Centre at Moorfields Eye Hospital NHS Foundation Trust, and UCL Institute of Ophthalmology, London, United Kingdom (grant no. BRC2_009). Additional support was provided by The Special Trustees of Moorfields Eye Hospital, London, United Kingdom (grant no. ST 12 09). Data analysis was carried out using the RAVEN computing cluster, maintained by the ARCCA group at Cardiff University ARCCA and the BLUE CRYSTAL3 computing cluster maintained by the HPC group at the University of Bristol.

## Author contributions

Conceptualization: AP, JAG. Funding Acquisition: CW, JAG. Formal Analysis: AP. Data Curation: AP, PH, JAG. Writing – Original Draft Preparation: All authors. Writing – Review & Editing: All authors.

